## Supplemental Figures for "BMP2 and BMP7 cooperate with H3.3K27M to promote quiescence and invasiveness in pediatric diffuse midline gliomas"

### Figure supplements

**Figure 1-figure supplement. BMP pathway activation coincides with high BMP2/7 levels in *ACVR1* WT-H3.3K27M pDMG**

**A.** Principal Component Analysis (PCA) of *ACVR1* WT (red) and mutant pDMG (blue) samples based on transcriptomic data from cohort 2.

**B.** Heatmap representing the transcriptomic expression levels of BMP target genes between *ACVR1* WT (red) and mutant pDMG (blue) in cohort 2. H3 mutational status is specified as WT (green), mutated on variant H3.1 (brown) or H3.3 (purple). Normalized and centered gene expression levels are color-coded with a blue (low expression) to red (high expression) gradient. Samples in columns are clustered using Euclidean distance. The BMP targets gene list is presented in Supplementary Table 1.

**C.** TGF-β/BMP pathway activity inferred from a specific genes-response signature using PROGENy algorithm in *ACVR1* WT (red) versus mutant pDMG (blue) (cohort 2). No significant difference was observed between both groups.

**D**. Comparison of the expression levels of BMP ligands, antagonists, receptors, and effectors between *ACVR1* WT (red) versus mutant pDMG (blue) in cohort 2. p-values are indicated for each gene.

**E.** Boxplot of BMP ligands expression (vst-normalized) in cohort 3.

**F.** Boxplot of BMP2 and 7 expressions (vst-normalized) in cohort 2, according to *ACVR1* and *H3* status.

**G.** Pattern of expression of BMP4 in developing brain. **Left panel:** heatmaps showing relative BMP4 expression by *in situ* hybridisation (ISH) in murine brain across development obtained from the ALLEN Developing Mouse Brain Atlas. Normalized and scaled gene expression levels are color-coded with a yellow (low expression) to red (high expression) gradient. Developmental stage is mentioned in rows with pre- and post-natal stage color-coded in dark and light blue, respectively. Different brain regions are indicated in columns as following: RSP: rostral secondary prosencephalon; Tel: telencephalic vesicle; PedHy: peduncular caudal hypothalamus; P3: prosomere 3; P2: prosomere 2; P1: prosomere 1; M: midbrain; PPH: prepontine hindbrain; PH: pontine hindbrain; PMH: pontomedullary hindbrain; MH: medullary hindbrain (medulla). **Right panel:** spatiotemporal gene expression data of BMP4 expression from human developing and adult brain samples obtained from the Human Brain Transcriptome[32]. The vertical line indicates birth at 266 days. Each curve represents a part of the brain as following: NCX: neocortex (dark blue); HIP: hippocampus (light blue); AMY: amygdala (orange); STR: striatum (black); MD: mediodorsal nucleus of the thalamus (green, in bold); CBC: cerebellar cortex (red).

**Figure 2-figure supplement. H3.3K27M mutant context potentiates BMP7-induced transcriptomic and phenotypic switch in a glioma model**

**A.** *BMP2*, *BMP7* and *BMP4* normalized expression from transcriptomic data of Res259-H3.3WT (green) *versus* Res259-H3.3K27M (purple). Means +/- std are represented (n = 3).

**B.** *BMP4* expression between Res259-H3.3WT (green) and H3.3K27M (purple). Gene expression was analysed by QRT-PCR relative to *HPRT1* expression. Means +/- std are represented (n = 3). ns: non-significant p-value.

**C**. *ID1* and *ID2* expressions between SF188- (upper panel) and Res259- (bottom panel) H3.3WT (green) and H3.3K27M (purple) following BMP7 treatment at indicated time points from 0 to 48 hr (x-axis). Means +/- std are represented (n = 3). *:$p < 0.05$, ns: non-significant.

**D**. Western-blot analysis of SMAD1/5/8 phosphorylation on Ser463/465 (SMAD1/5) and Ser465/467 (SMAD8) (pSMAD1/5/8) levels in SF188- and Res259-WT and H3.3K27M upon BMP7 treatment. Total SMAD1 and β-actin are used as loading control for the indicated conditions (n = 3). A representative experiment is shown.

**E**. Venn diagram showing the number of differentially expressed genes (DEG) and the corresponding percentages compared to all DEG in each condition: Res259-H3.3WT versus Res259-H3.3K27M without BMP7 treatment (grey), Res259-H3.3WT versus Res259-H3.3WT treated with BMP7 for 3h (green), Res259-H3.3K27M versus Res259-H3.3K27M treated with BMP7 for 3hr (purple).

**F**. Heatmap representing the transcriptomic expression levels of the 29 DEG specifically in Res259-H3.3K27M versus Res259-H3.3K27M treated with BMP7 for 3 hr. Samples are placed in columns such as Res259-H3.3WT (green) and -H3.3K27M (purple), control (light blue) or 3 hr BMP7 treatment (blue). Normalized and centered gene expression levels are color-coded with a blue (low expression) to red (high expression) gradient (n = 3).

**G**. Volcano plots of DEG in the E2F target signature between Res259-H3.3WT control *versus* BMP7-treated (left panel) and Res259-H3.3K27M control *versus* BMP7-treated (right panel). Fold change is indicated on the x-axis and statistical significance on the y-axis. Black dots: non-differentially expressed. Red dots: differentially expressed.

**H**. Viable cell count of SF188- and Res259-H3.3WT and H3.3K27M after 72 hr of BMP7 treatment (dark blue), normalized to non-treated condition (light blue). Means +/- std are represented (n = 3). *: $p < 0.05$, ns: non-significant.

**I**. Senescence-associated beta-galactosidase activity (SA-β-gal) assessment experiment on Res259-H3.3WT and Res259-H3.3K27M control or treated with BMP7 for 72 h with SA-β-gal staining. Right panel: representative microscopic photos. Arrows indicate SA-β-gal+ cells. Left panel: quantification of cells positive for SA-β-gal activity. Means +/- std are represented (n = 3). *:p < 0.05, ns: non-significant.

**J**. Decisional tree algorithm to study the specific impact of potentiation of H3.3K27M by BMP7 (left) and the synergy of both (right). DE: differentially expressed. DEG: differentially expressed genes. FC: fold-change. K27M: H3.3K27M mutation. WT : H3.3wt. For the last step of decisional tree 2 (right panel), genes are selected such as log2(FC(K27M+BMP7)/(K27M)) > 1.5 x log2(FC(WT+BMP7)/(WT)).

**Figure 3-figure supplement. Combined tumor-autonomous BMP2/BMP7 expression drives a quiescent-invasive tumor cell state in pDMG**

**A.** BMP ligands expression analysed by QRT-PCR relative to 5 housekeeping genes levels in three different DIPG cell lines. HSJD-DIPG-007: *ACVR1* mutant (grey). HSJD-DIPG-012: *ACVR1* WT (dark blue). HSJD-DIPG-014: *ACVR1* WT (light blue). Means +/- std are represented (n = 2).

**B.** Western-blot analysis of pSMAD1/5/8, total SMAD1/5/8 on HSJD-DIPG-012 spheroids, treated or not with BMP2. GAPDH was used as a loading control. A representative experiment out of 3 is shown.

**C.** Growth monitoring of HSJD-DIPG-014 (top panel) and HSJD-DIPG-013 (bottom panel) following recombinant BMP2 treatment. Means +/- std are represented. *: p<0.05, **: p<0.01, ***: p<0.001.

**D.** Growth monitoring of BT245 and DIPGXIII, before and after ($KO^{K27M}$) edition of the H3.3K27M mutation. Representative pictures of BT245 are shown on the left. Means +/- std are represented on the right.

**E.** Impact of BMP2 or LDN treatment on tumor cells invasion. **Left panel:** representative images of HSJD-DIPG-013 spheroids embedded in Matrigel, after 48 hr of BMP2 or LDN-193189 treatment. Scale bar = 250 µm. **Right panel:** Invasion was quantified as the mean value of four independent experiments and represented as a graph. *: p<0.05. BMP2: 10ng/mL. LDN:1 µM.

**F.** Violin plots of BMP receptors in two (P-1764_S-1766 and P-3407_S-3447) of the 10 samples based on the TGF-β/BMP score. The TGF-β/BMP-low group is colored in light blue and the TGF-β/BMP-high group is colored in blue.

**G.** Dotplot of the first 10 significantly enriched pathways (FDR <= 0.05) in TGF-β/BMP-low cells for each of the 10 *ACVR1* WT-H3.3K27M pDMG, ranked by number of samples with a significant enrichment, using the GO Biological Process database. Only dots of significant enrichments are shown for each sample. Dot color represents the -log10(p-value) and ranges from blue (high p-value) to yellow (low p-values). Dot size is proportional to the overlap of DE genes and the genes of a geneset.

**H.** Score of the pDMG niche signatures identified by [Ren et al., 2023], namely: “Tumor core”, “Vascular niche”, “Invasive niche” and “Hypoxic niche” in each of the 3 Visium samples. Color ranges from blue (low score value) to red (high score value).

**I.** Scatter plots correlating PROGENy TGF-β/BMP pathway activity with the "Invasive niche" score for each Visium spot of Sample-2 and Sample-3. The correlation coefficient was computed using Pearson's method. ****: p-value < 2.e-16.

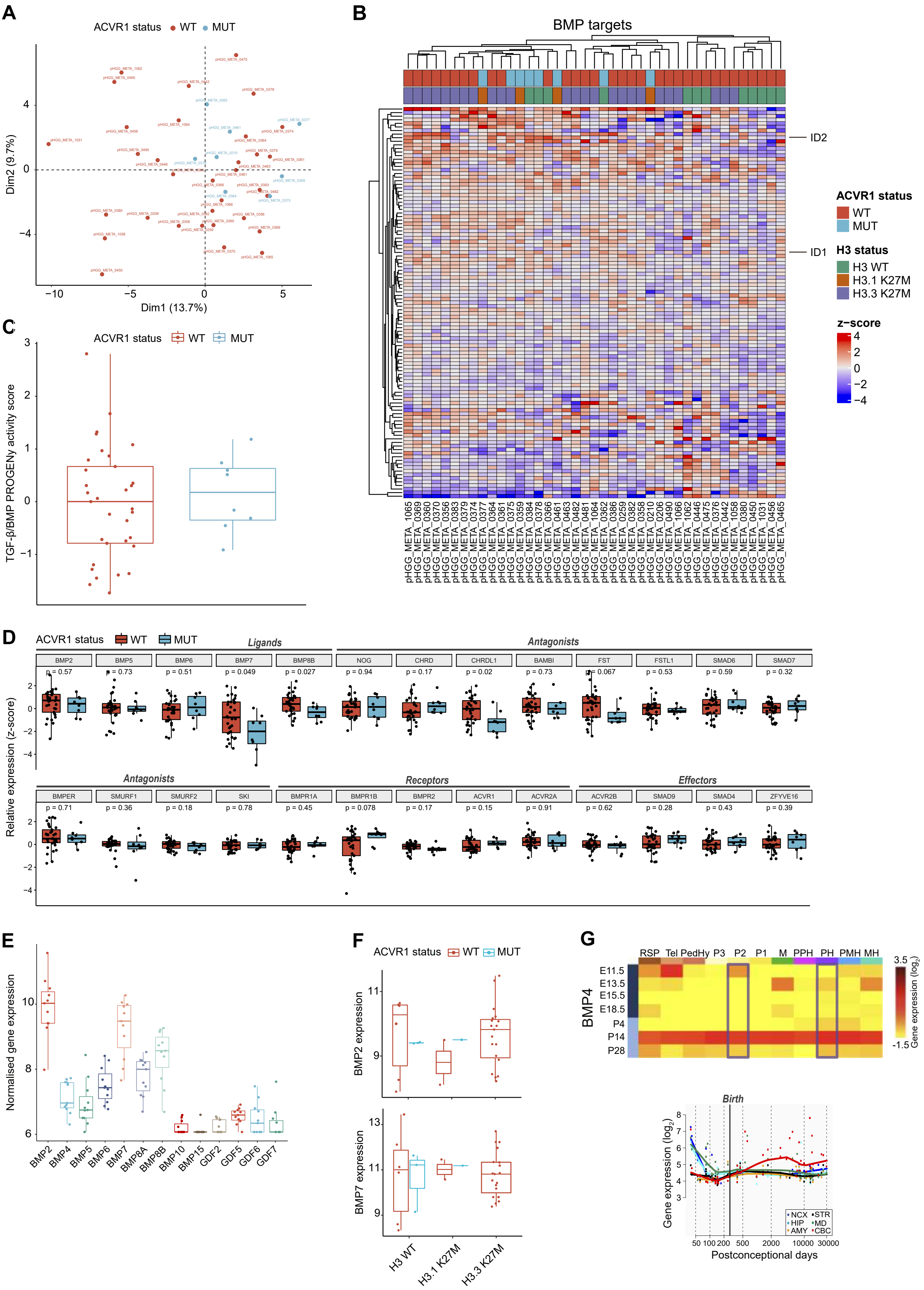

A
ACVR1 status WT MUT
Dim1 (13.7%)
Dim2 (9.7%)
B
BMP targets
ID2
ID1
ACVR1 status
WT
MUT
H3 status
H3 WT
H3.1 K27M
H3.3 K27M
z−score
C
ACVR1 status WT MUT
TGF-β/BMP PROGENy activity score
D
ACVR1 status WT MUT
Ligands
Antagonists
Receptors
Effectors
BMP2 BMP5 BMP6 BMP7 BMP8B NOG CHRD CHRDL1 BAMBI FST FSTL1 SMAD6 SMAD7
p = 0.57 p = 0.73 p = 0.51 p = 0.049 p = 0.027 p = 0.94 p = 0.17 p = 0.02 p = 0.73 p = 0.067 p = 0.53 p = 0.59 p = 0.32
BMPER SMURF1 SMURF2 SKI BMPR1A BMPR1B BMPR2 ACVR1 ACVR2A ACVR2B SMAD9 SMAD4 ZFYVE16
p = 0.71 p = 0.36 p = 0.18 p = 0.78 p = 0.45 p = 0.078 p = 0.17 p = 0.15 p = 0.91 p = 0.62 p = 0.28 p = 0.43 p = 0.39
Relative expression (z-score)
E
Normalised gene expression
BMP2 BMP4 BMP5 BMP6 BMP7 BMP8A BMP8B BMP10 BMP15 GDF2 GDF5 GDF6 GDF7
F
ACVR1 status WT MUT
BMP2 expression
BMP7 expression
H3 WT H3.1 K27M H3.3 K27M
G
RSP Tel PedHy P3 P2 P1 M PPH PH PMH MH
BMP4
E11.5 E13.5 E15.5 E18.5 P4 P14 P28
Gene expression (log2)
Birth
Gene expression (log2)
Postconceptional days
NCX STR HIP MD AMY CBC

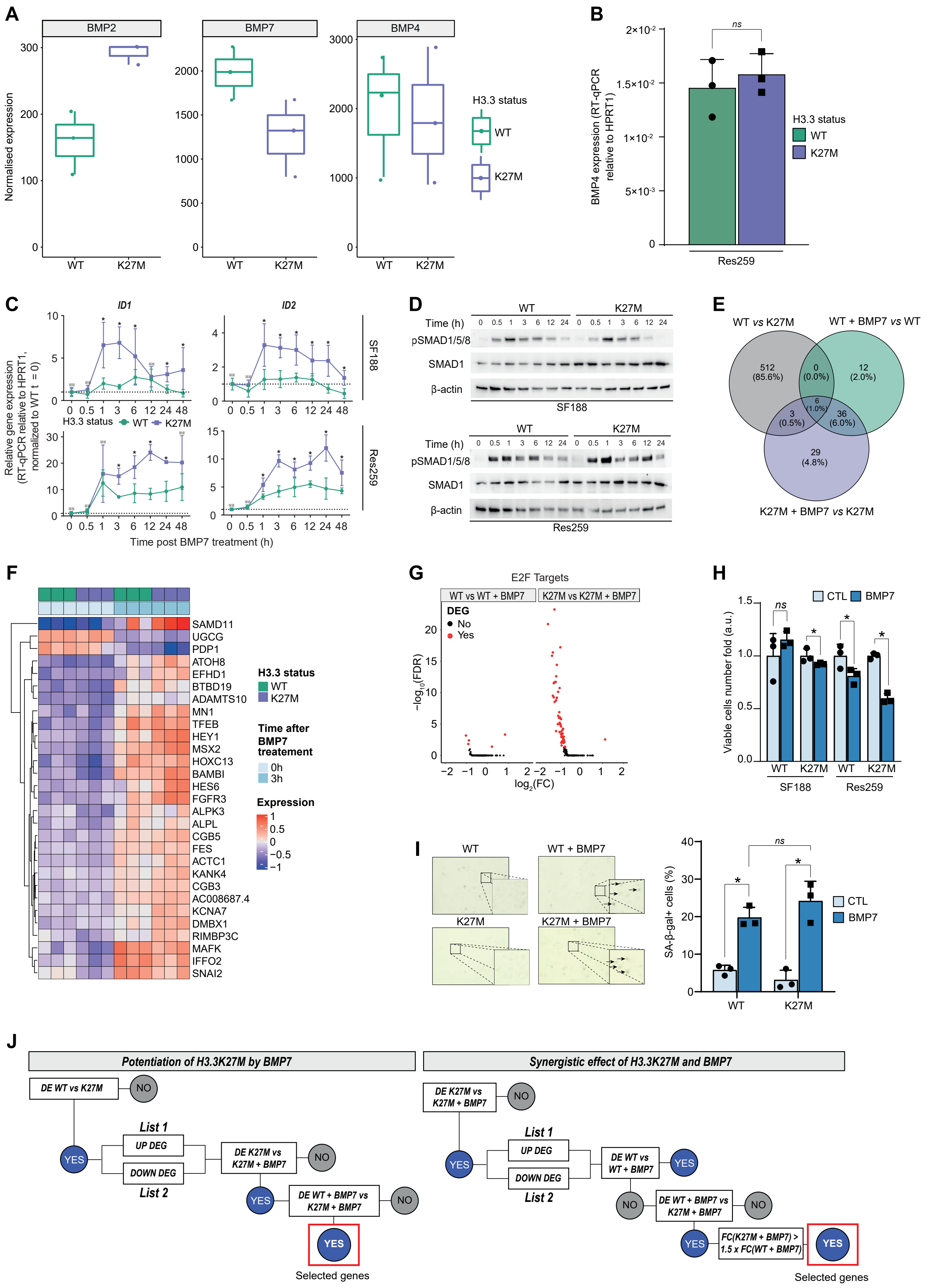

A
BMP2
BMP7
BMP4
Normalised expression
WT
K27M
H3.3 status
WT
K27M
B
BMP4 expression (RT-qPCR relative to HPRT1)
ns
H3.3 status
WT
K27M
Res259
C
ID1
ID2
Relative gene expression (RT-qPCR relative to HPRT1, normalized to WT t = 0)
H3.3 status WT K27M
SF188
Res259
Time post BMP7 treatment (h)
D
Time (h)
WT
K27M
pSMAD1/5/8
SMAD1
β-actin
SF188
Res259
E
WT vs K27M
WT + BMP7 vs WT
K27M + BMP7 vs K27M
512 (85.6%)
0 (0.0%)
12 (2.0%)
6 (1.0%)
3 (0.5%)
36 (6.0%)
29 (4.8%)
F
SAMD11
UGCG
PDP1
ATOH8
EFHD1
BTBD19
ADAMTS10
MN1
TFEB
HEY1
MSX2
HOXC13
BAMBI
HES6
FGFR3
ALPK3
ALPL
CGB5
FES
ACTC1
KANK4
CGB3
AC008687.4
KCNA7
DMBX1
RIMBP3C
MAFK
IFFO2
SNAI2
H3.3 status
WT
K27M
Time after BMP7 treatement
0h
3h
Expression
G
E2F Targets
WT vs WT + BMP7
K27M vs K27M + BMP7
DEG
No
Yes
−log10(FDR)
log2(FC)
H
CTL
BMP7
Viable cells number fold (a.u.)
ns
WT
K27M
SF188
Res259
I
WT
WT + BMP7
K27M
K27M + BMP7
SA-β-gal+ cells (%)
ns
CTL
BMP7
J
Potentiation of H3.3K27M by BMP7
DE WT vs K27M
NO
YES
List 1
UP DEG
DOWN DEG
List 2
DE K27M vs K27M + BMP7
DE WT + BMP7 vs K27M + BMP7
Selected genes
Synergistic effect of H3.3K27M and BMP7
DE K27M vs K27M + BMP7
DE WT vs WT + BMP7
DE WT + BMP7 vs K27M + BMP7
FC(K27M + BMP7) > 1.5 x FC(WT + BMP7)
Selected genes

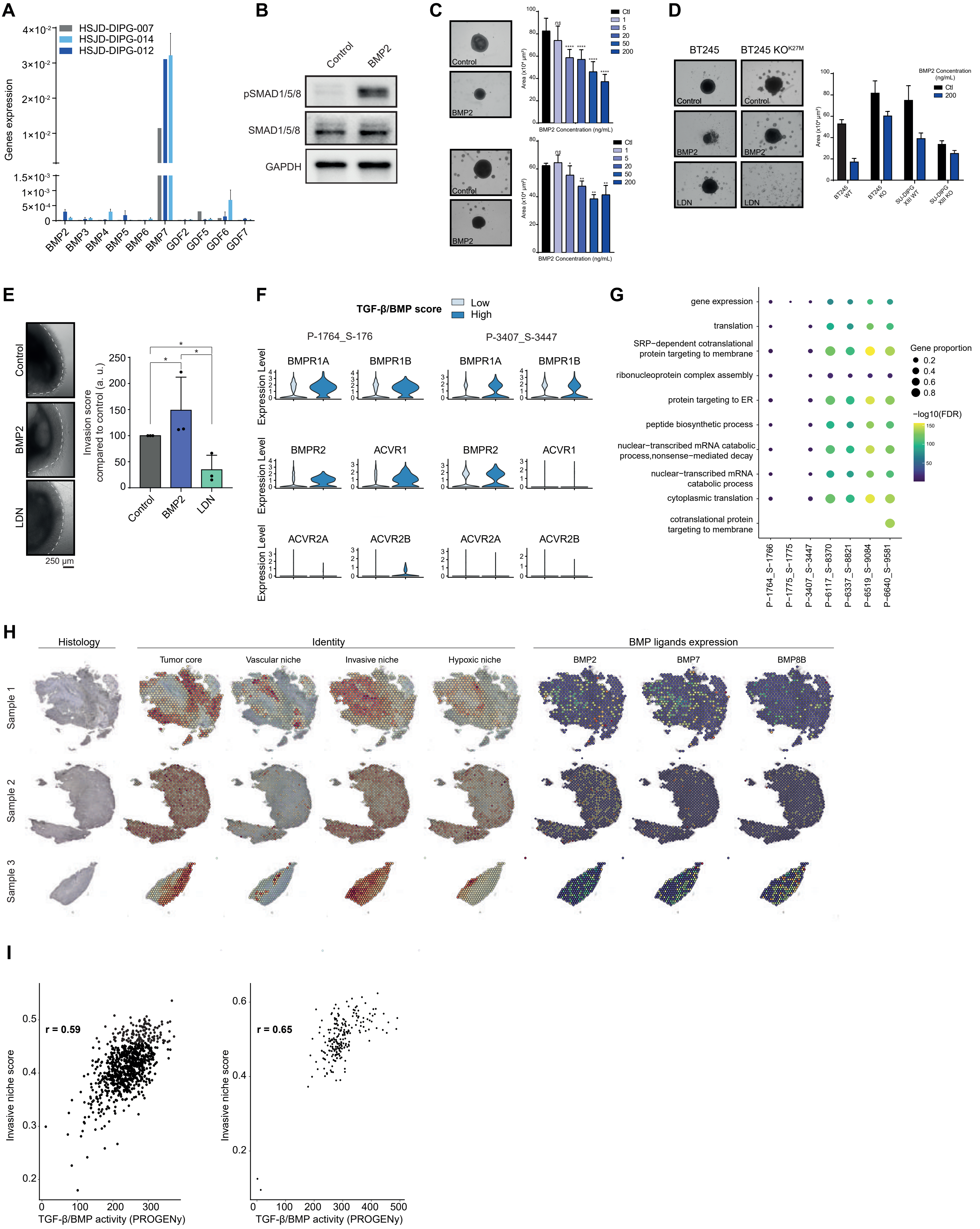
A
HSJD-DIPG-007
HSJD-DIPG-014
HSJD-DIPG-012
Genes expression
BMP2 BMP3 BMP4 BMP5 BMP6 BMP7 GDF2 GDF5 GDF6 GDF7
B
Control
BMP2
pSMAD1/5/8
SMAD1/5/8
GAPDH
C
Control
BMP2
BMP2 Concentration (ng/mL)
Ctl
D
BT245
BT245 KO
Control
BMP2
LDN
BMP2 Concentration (ng/mL)
BT245 WT
BT245 KO
SU-DIPG XIII WT
SU-DIPG XIII KO
E
Control
BMP2
LDN
250 µm
Invasion score compared to control (a. u.)
F
TGF-β/BMP score
Low
High
P-1764_S-176
P-3407_S-3447
Expression Level
BMPR1A
BMPR1B
BMPR2
ACVR1
ACVR2A
ACVR2B
G
gene expression
translation
SRP−dependent cotranslational protein targeting to membrane
ribonucleoprotein complex assembly
protein targeting to ER
peptide biosynthetic process
nuclear−transcribed mRNA catabolic process,nonsense−mediated decay
nuclear−transcribed mRNA catabolic process
cytoplasmic translation
cotranslational protein targeting to membrane
Gene proportion
−log10(FDR)
P−1764_S−1766
P−1775_S−1775
P−3407_S−3447
P−6117_S−8370
P−6337_S−8821
P−6519_S−9084
P−6640_S−9581
H
Histology
Identity
BMP ligands expression
Tumor core
Vascular niche
Invasive niche
Hypoxic niche
BMP2
BMP7
BMP8B
Sample 1
Sample 2
Sample 3
I
r = 0.59
r = 0.65
Invasive niche score
TGF-β/BMP activity (PROGENy)
